# Identification and Implications for Tumor Heterogeneity of a DNA Methylation-Based Signature Classifying Pancreatic Ductal Adenocarcinoma Based on their Cellular Origin

**DOI:** 10.1101/2025.03.07.642043

**Authors:** Ilianna Zoi, Memoona Rajput, Adrián Holguín-Horcajo, Malak Haidar, Davide Brusa, Ken Chu, Jane Xie, Mario Shields, Sofia Morgadinho Ferreira, Ben Stanger, Laura D. Attardi, Janel Kopp, Marc P. Stemmler, Rémy Nicolle, Meritxell Rovira, Patrick Jacquemin

**Affiliations:** Université catholique de Louvain, de Duve Institute, Brussels, Belgium; Pancreas Regeneration: Pancreatic Progenitors and Their Niche Group, Regenerative Medicine Program, Institut d’Investigació Biomèdica de Bellvitge [IDIBELL, L’Hospitalet de Llobregat, Spain; Department of Physiological Science, School of Medicine, Universitat de Barcelona, L’Hospitalet de Llobregat, Spain; CytoFlux-IREC Flow Cytometry Platform, Université catholique de Louvain, Institute for Experimental and Clinical Research, Brussels, Belgium; Department of Cellular and Physiological Sciences, Life Sciences Institute, University of British Columbia, Vancouver, Canada; Department of Medicine, Perelman School of Medicine, University of Pennsylvania, Philadelphia, Pennsylvania; Department of Pathology and Stony Brook Cancer Center, Renaissance School of Medicine, Stony Brook University, Stony Brook, New York; Departments of Radiation Oncology and Genetics, Stanford University School of Medicine, Stanford, California; Department of Experimental Medicine 1, Nikolaus-Fiebiger Center for Molecular Medicine, FAU University Erlangen-Nürnberg, Erlangen, Germany; Université Paris Cité, Centre de Recherche Sur L’Inflammation (CRI), INSERM, U1149, Paris, France

## Abstract

**Background:** Pancreatic ductal adenocarcinoma (PDAC) arises from distinct cellular origins, yet the extent to which DNA methylation patterns from normal pancreatic cells are preserved in tumor cells remains unclear. Identifying cell-of-origin signatures may enhance PDAC classification and therapeutic stratification.

**Objective:** To determine whether DNA methylation signatures in normal acinar and ductal pancreatic cells are retained in PDAC cell lines and to develop a robust classifier for distinguishing tumor origins.

**Design:** We performed DNA methylation profiling using the Illumina Infinium Mouse MethylationEPIC array on normal acinar and ductal cells and their PDAC derivatives in genetically engineered mouse models (GEMMs). Differential methylation analysis, and hierarchical clustering were used to identify and validate a conserved cell-of-origin DNA methylation signature. A logistic regression model was developed for classification.

**Results:** We identified 178 CpG sites that remain preserved during tumorigenesis and effectively distinguished acinar- and ductal-derived PDAC cell lines. This signature was validated across independent sample sets, primary tumors, and orthotopic allografts. It successfully classified cell lines of unknown origin, including PDAC samples from KPC mice, and revealed the impact of oncogenic mutations on tumor fate. A logistic regression model supported these findings, confirming the robustness of the classification approach. Furthermore, the cell of origin influenced key PDAC characteristics, including treatment response, highlighting its potential role in molecular subtyping and patient stratification.

**Conclusion:** A preserved DNA methylation signature during pancreatic carcinogenesis distinguishes PDAC origins and influences tumor behavior. These results highlight the potential of DNA methylation profiling for tumor classification and personalized treatment strategies. They also raise important questions about the relevance of KPC mice as a preclinical model and the mechanisms driving PDAC heterogeneity.

**What is already known on this topic:**

- Human PDAC exhibits significant heterogeneity, with molecular subtyping (classical vs basal-like) providing some insights into tumor behavior and clinical outcomes.
- Mouse acinar and ductal cells can give rise to PDAC, influencing tumor characteristics and survival outcomes.
- DNA methylation is a powerful tool for tracing cellular identity and distinguishing cancer subtypes based on epigenetic profiles.

**What this study adds:**

- A cell-of-origin methylation signature is preserved during mouse carcinogenesis and across diverse experimental settings, providing a reliable tool for tumor classification.
- A DNA methylation-based classification system reliably distinguishes between acinar- and ductal-origin PDAC, filling a critical gap in methods to trace tumor lineage.
- Acinar- and ductal-derived PDACs exhibit distinct methylation patterns that correlate with differences in tumor behavior, such as chemoresistance, highlighting the biological relevance of cellular origin in PDAC.
- The study provides new insights into how cell-of-origin influences PDAC heterogeneity and could lead to more precise therapeutic strategies tailored to the tumor’s lineage.

**How this study might affect research, practice or policy:**

- This study provides a new, reliable method for classifying PDAC based on its cellular origin, which could significantly improve tumor classification in both preclinical and clinical settings, aiding in more accurate diagnoses and prognostic predictions.
- The identification of distinct methylation patterns linked to tumor behavior offers valuable insights for developing personalized treatment strategies, as therapies could be tailored based on the tumor’s cellular origin and associated molecular characteristics.
- The preservation of cell-of-origin methylation signature suggests the potential for developing a universal biomarker for PDAC classification, which could guide future clinical trials, therapeutic targeting, and patient stratification in PDAC care.

## Introduction

Pancreatic ductal adenocarcinoma (PDAC) is among the most lethal cancers and is projected to become the second leading cause of cancer-related deaths by 2030 [1]. The aggressive nature of PDAC, coupled with late-stage diagnosis, contributes to its poor prognosis. PDAC is characterized by significant intertumour and intratumour heterogeneity, which complicates treatment and diagnosis [2,3]. Molecular subtyping has been essential in understanding this heterogeneity. PDAC is mainly classified into classical and basal-like subtypes, providing insights into distinct clinical outcomes [4–6]. The classical subtype is typically associated with a more differentiated phenotype and better prognosis, while the basal-like subtype is more aggressive and poorly differentiated [5].

The heterogeneity of PDAC extends to its cellular origins. Genetically engineered mouse models (GEMMs) have been pivotal in studying the cell of origin of pancreatic cancer, demonstrating that both acinar and ductal cells can give rise to PDAC. This duality in cellular origin is believed to influence the pathophysiological and molecular characteristics of the resulting tumors [7–10]. Despite the typical ductal histology of pancreatic tumors, research has shown that acinar cells, responsible for secreting digestive enzymes, can undergo acinar-to-ductal metaplasia (ADM) under inflammatory stress, and subsequently progress to pancreatic intraepithelial neoplasia (PanIN) [11], leading ultimately to PDAC. On the other hand, ductal cells, which line the pancreatic ducts, can directly give rise to PDAC, bypassing the intermediate neoplastic precursor lesions [7,8,10]. Acinar-derived PDACs have been associated with the classical subtype, while ductal-derived PDACs correlate with the basal-like subtype [10,12,13]. Mouse pancreatic tumors with ductal origin have been further linked with immunosuppression [13] and lower survival rates [7–9]. In addition to these results obtained in mice, strong evidence was provided that human PDACs can also originate from both acinar and ductal cells [14].

DNA methylation has emerged as a powerful tool for understanding the epigenetic landscape of tumors. Studies have shown that distinct DNA methylation patterns characterize each stage in the transformation process of a pancreatic cancer cell, from normal to premalignant and from malignant to invasive states [15,16]. Moreover, DNA methylation reflects cellular identity, providing an excellent means to trace the origins of cancer cells [17,18]. Building on this concept, a recent study identified two distinct PDAC subtypes with differential DNA methylation patterns, suggesting potential links to acinar and ductal cellular identities [12]. However, their study left unresolved whether these findings could be used to develop a practical method for classifying PDAC samples based on their cell of origin.

In this context, our study addresses this critical gap by performing mouse methylation profile analyses on FACS-purified normal acinar and ductal cells, and acinar-derived and ductal-derived PDAC cell lines, to identify maintained DNA methylation footprints corresponding to the two major cell origins of PDAC. The identified cell-of-origin signature was validated by analyzing additional known-origin PDAC cell lines, as well as primary tumors and allografts. A logistic regression model was then employed to classify PDAC cell lines of unknown origin based on this signature. Remarkably, PDAC cell lines classified as acinar-origin were enriched in immune response pathways and showed greater response to chemotherapy agents. These findings establish a cell-of-origin-based classification framework, suggest potential therapeutic implications, and question the established mechanisms explaining PDAC heterogeneity.

## Materials and Methods

Detailed descriptions of all materials and experimental methods can be found in online supplementary methods.

## Results

### Preserved Cell-of-Origin DNA Methylation Signatures in Normal Pancreatic Cells and PDAC Cell Lines

To determine the DNA methylation profiles of normal pancreatic cells and PDAC cell lines, we performed Illumina Infinium Mouse MethylationEPIC array BeadChip (285K probes) on 12 mouse samples (discovery group) (Supplementary Table S1). We first used GEMMs to trace and purify normal acinar and ductal cells. Acinar cells were traced using *ElastaseCreER; ROSA26^Yellow^ ^Fluoresence^ ^Protein^ ^(YFP)^* (Acinar:YFP) mice, which express YFP in acinar cells upon tamoxifen treatment. Ductal cells were traced using *Sox9eGFP* (Ductal:EGFP) mice, in which Sox9-expressing ductal cells are labeled with EGFP. To ensure purity, we FACS-isolated acinar, YFP+ (Ela+; n = 3) and ductal, EpCAM+Sox9eGFP+SSC^low^(Sox9+; n = 3) cells (Supplementary Figure 1A, [19]).

Principal component analysis (PCA) of the global DNA methylation levels (beta values) of CpG sites revealed distinct methylation patterns between acinar and ductal cells, with PC1 accounting for 70% of variance (Supplementary Figure 1B). Differential methylation analysis identified 2,346 differentially methylated probes (DMPs) between acinar and ductal cells (normal DMPs), with an absolute mean difference (Δβ) of at least 0.35 and an FDR adjusted p-value less than 0.05 (Figure 1A, B). These normal DMPs were primarily located in open sea regions (86.95%), with 82.1% hypermethylated in acinar cells and 17.9% in ductal cells (Supplementary Figure 1C).

**Figure 1.**
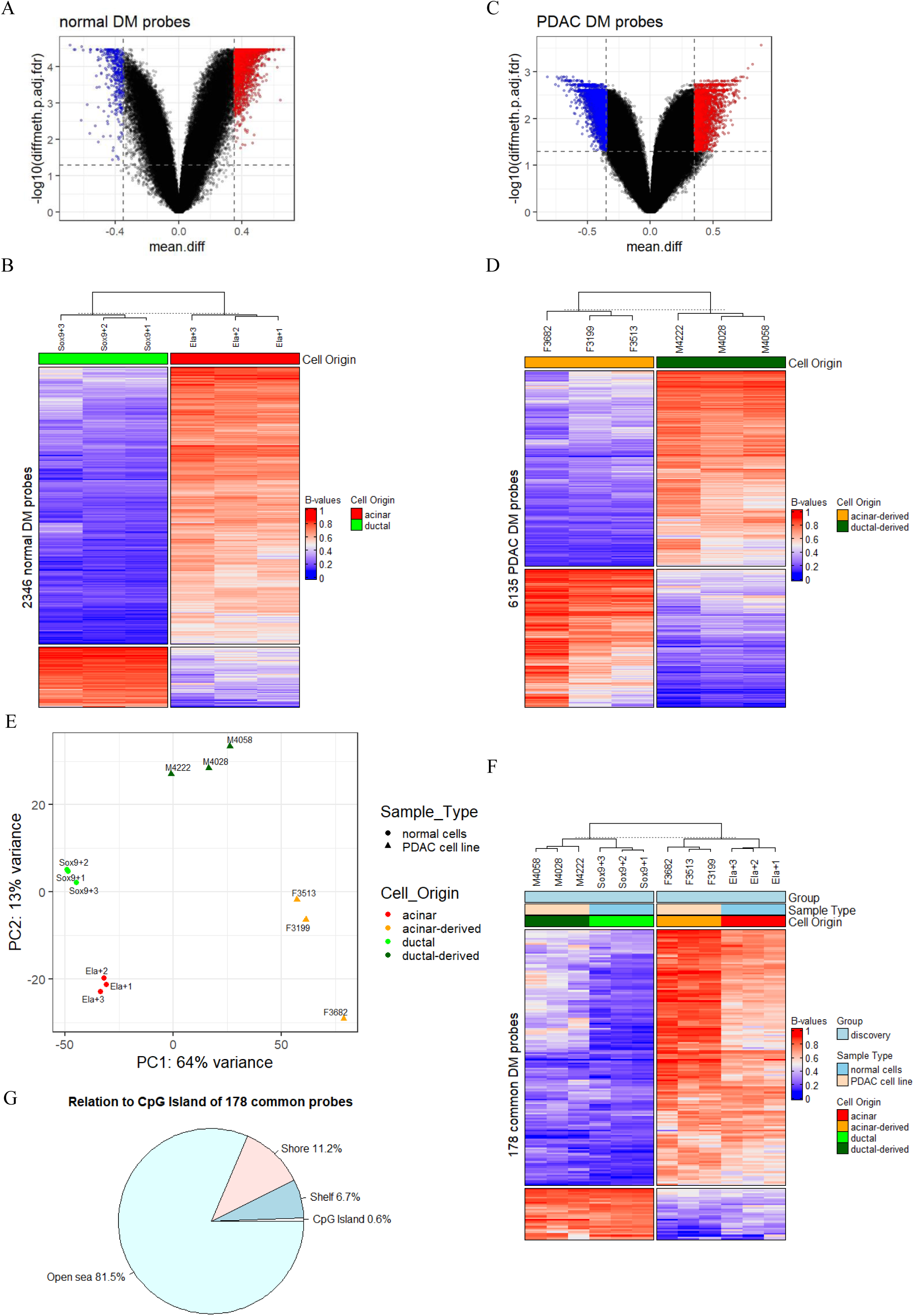
DNA methylation analysis reveals distinct epigenetic signatures between acinar and ductal cells, their corresponding PDAC cell lines, and the identification of lineage-specific CpG sites for PDAC classification. (A) Volcano plots of DMPs in acinar vs ductal cells. Vertical dashed lines indicate an absolute mean difference of 0.35 and horizontal line corresponds to an FDR adjusted p-value equal to 0.05. Red dots represent hypermethylated probes and blue dots represent hypomethylated probes for each comparison. (B) Hierarchical clustering of normal cells using the 2,346 DMPs from the acinar vs ductal comparison. (C) Volcano plots of DMPs in acinar-derived (Acinar1:KP) vs ductal-derived (Ductal1:KP) cell lines. Vertical dashed lines indicate an absolute mean difference of 0.35 and horizontal line corresponds to an FDR adjusted p-value equal to 0.05. Red dots represent hypermethylated probes and blue dots represent hypomethylated probes for each comparison. (D) Hierarchical clustering of PDAC cell lines using the 6,135 DMPs from the acinar-derived vs ductal-derived comparison. (E) Principal component analysis (PCA) of the beta values from normal FACS-purified cells and PDAC cell lines. Percentages indicate the proportion of variance explained by each component. Acinar (Acinar:YPF): n = 3; Ductal (Ductal:EGFP): n = 3; Acinar-derived (Acinar1:KP): n = 3; Ductal-derived (Ductal1:KP): n = 3. (F) Hierarchical clustering of the discovery samples (normal cells and PDAC cell lines) using the 178 common probes found in normal and PDAC DMPs. (G) Relation to CpG island of the 178 common probes. CpG island are DNA regions frequently found in gene promoters and rich in CpG sites, compared to the rest of the genome. Based on their proximity to CpG islands, other DNA regions are categorized as shores, shelves, and open sea, with shores being the closest, and the open sea the most distant from CpG islands.

Next, we generated acinar-derived and ductal-derived PDAC mouse models. Acinar-derived PDAC samples were obtained from ElaCreER; LSL-Kras^G12D/+^; LSL-Trp53^R172H/+^ or ElaCreER; LSL-Kras^G12D/4AKO^; LSL-Trp53^R172H/+^ (Acinar1:KP) mice after tamoxifen treatment and the induction of chronic inflammation through long-term cerulein treatment. Ductal-derived tumors were formed in Sox9CreER; LSL-Kras^G12D/+^; LSL-Trp53^f/f^ (Ductal1:KP) mice after tamoxifen treatment. To minimize confounding effects from stromal cells in PDAC tumors, we derived cell lines from acinar-derived (n = 3) and ductal-derived (n = 3) tumors. PCA of all methylation sites showed clear separation of acinar- and ductal-derived PDAC cell lines along PC1 (64% variance), reflecting their cellular origins (Supplementary Figure 1D). Differential methylation analysis revealed 6,135 DMPs between acinar-derived and ductal-derived PDAC cell lines (PDAC DMPs), with an |Δβ| of ≥ 0.35 and an FDR adjusted p-value less than 0.05 (Figure 1C). These PDAC DMPs were predominantly found outside CpG islands (open sea regions: 62.8%), with 58.6% more methylated in ductal-derived cancer cells (Supplementary Figure 1E). Hierarchical clustering based on PDAC DMPs showed the clear segregation of PDAC cell lines by their cellular origins (Figure 1D). The integration of normal and PDAC cell methylomes showed that cellular transformation was the primary driver of DNA methylation variation (PC1: 64% variance; Figure 1E). Specifically, among the 1,925 probes hypermethylated in acinar cells, 614 (31.9%) displayed reduced methylation in acinar-derived PDAC lines, whereas 1,028 (49.4%) gained methylation in ductal-derived PDAC lines (Supplementary Figure 2A). Likewise, out of the 421 probes methylated in ductal cells, 48 (11.4%) showed reduced methylation in ductal-derived PDAC lines, while 297 (70.5%) became more methylated in acinar-derived PDAC lines (Supplementary Figure 2A). Despite these changes, certain lineage-specific gene markers retained their promoter methylation profiles (Supplementary Figure 2B, C).

To identify CpG sites that maintain their methylation status through pancreatic carcinogenesis and reflect the cell of origin, we intersected normal and PDAC DMPs, identifying 178 common probes (Supplementary Figure 2D, Supplementary Table S2). Hierarchical clustering using these 178 common probes effectively separated acinar-origin and ductal-origin samples, regardless of their transformation state (Figure 1F). These probes were mainly located in open sea regions (81.5%) (Figure 1G), with 148 hypermethylated probes in acinar-origin samples and 30 hypermethylated probes in ductal-origin samples. These common probes constitute the so-called cell-of-origin signature.

Collectively, these analyses show the presence of distinct methylation patterns in mouse acinar and ductal cells and their PDAC derivatives, identifing a cell-of-origin DNA methylation signature that is preserved during pancreatic carcinogenesis.

### Validation of the Cell-of-Origin DNA Methylation Signature

To validate our cell-of-origin signature, we profiled the methylome of six additional cell lines derived from acinar-derived and ductal-derived tumors in our different laboratories (validation cell lines) (Supplementary Table S1). Acinar-derived tumors were induced in Ptf1a^CreER^; LSL- Kras^G12D/+^; LSL- Trp53^f/f^ (Acinar2:KP) mice (n = 3) [9]. Ductal-derived tumors were formed from the same model as previously described (Sox9CreER; LSL- Kras^G12D/+^; LSL- Trp53^f/f^ (Ductal2:KP) mice (n = 3) [9,10]. The addition of the six new PDAC cell lines in the PCA maintained a clear separation between normal and PDAC samples (PC1: 49% variance; Figure 2A). Furthermore, the PCA revealed a separation of the PDAC cell lines into two distinct groups (Figure 2A), with Acinar2:KP cells exhibiting a methylation profile more similar to Ductal1:KP and Ductal2:KP than Acinar1:KP cells (Figure 2B, Supplementary Figure 3A-D). This difference between Acinar1:KP and Acinar2:KP is likely due to the different approaches used to generate the two CreER transgenic lines (see Discussion).

**Figure 2.**
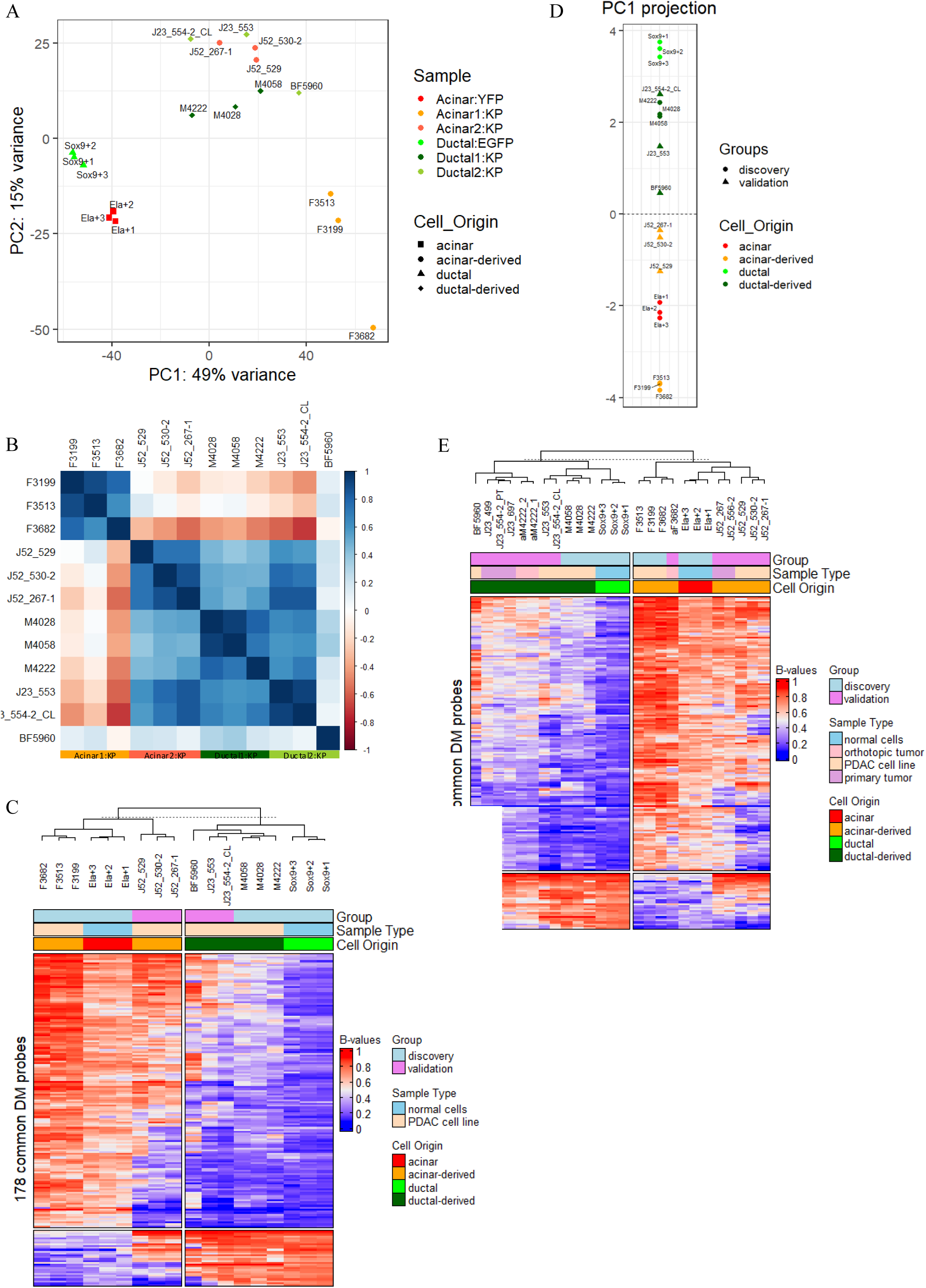
Analysis of PDAC samples with known origin confirms the reliability of the cell-of-origin DNA methylation signature for distinguishing acinar- and ductal-derived tumors. (A) PCA of normal acinar and ductal cells, and PDAC cell lines with known cell origin (discovery and validation lines) using all surveyed methylation sites. Percentages indicate the proportion of variance explained by each component. Acinar (Acinar:YPF): n = 3; Ductal (Ductal:EGFP): n = 3; Discovery acinar-derived (Acinar1:KP): n = 3; Discovery ductal-derived (Ductal1:KP): n = 3; Validation acinar-derived (Acinar2:KP): n = 3; Validation ductal-derived (Ductal2:KP): n = 3. (B) Pearson correlation plot of the methylation profiles of the discovery (Acinar1:KP: F3199, F3513, F3682, Ductal1:KP: M4028, M4058, M4222) and validation (Acinar2:KP: J52_529, J52_530-2, J52_267-1, Ductal2:KP: J23_553, J23_554-2_CL, BF5960) PDAC cell lines by the 10,000 most variable probes. (C) Hierarchical clustering of the discovery samples and the validation cell lines using the 178 common probes. (D) Projection of the validation cell lines onto the PC1 derived from the discovery samples using the cell-of-origin signature. (E) Hierarchical clustering of the discovery and validation samples using the cell-of-origin signature.

Despite this, hierarchical clustering using the cell-of-origin signature successfully distinguished acinar-origin from ductal-origin samples, with Acinar2:KP clustering with Acinar1:KP cells (Figure 2C). To further validate the 178 common probes, we projected the validation cell lines onto the PCA space derived from the discovery dataset (Supplementary Figure 4A). This allowed us to observe the clear separation between acinar-origin and ductal-origin samples (Figure 2D, Supplementary Figure 4B). These results support the robustness and reliability of our cell-of-origin DNA methylation signature across independent sample sets.

To assess the applicability of the signature to tumor tissues, we analyzed the DNA methylation of primary tumors from Acinar2:KP and Ductal2:KP mice, and allografts generated by orthotopically grafting Acinar1:KP and Ductal1:KP cells into NOD scid gamma (NSG) mice, eliminating immune cell interference (validation tumors). Hierarchical clustering using the signature probes accurately separated the validation tumors based on their cellular origin, despite the presence of abundant stroma (Figure 2E, Supplementary Figure 4C). By projecting the validation tumors onto the PCA space defined from the discovery dataset using the 178 common probes (Supplementary Figure 4A), we revealed distinct clusters for acinar- origin and ductal-origin samples (Supplementary Figure 4D). This further demonstrates the stability of the cell-of-origin DNA methylation signature across different PDAC models, highlighting its potential to distinguish the origin of tumors in various experimental settings.

### Clustering of Unknown-Origin Samples Using the Cell-of-Origin DNA Methylation Signature

To assess the effectiveness of the cell-of-origin signature in segregating samples with unknown cell origins (test group), we analyzed six cell lines with a similar mutational landscape to the previously studied cell lines. These cell lines were derived from Progenitor:KP (Pdx1Cre or Ptf1a^Cre^; LSL- Kras^G12D/+^; LSL- Trp53^R172H/+^) mice [20–22], where Cre recombinase is expressed in multipotent pancreatic progenitor cells during embryogenesis, leading to tumors potentially of acinar or ductal origin. Hierarchical clustering using the cell-of-origin signature revealed two distinct groups among the Progenitor:KP cell lines. Three cell lines (KPC792, KPC865, KPC-T) clustered with acinar-origin samples, while the remaining three (KPC827, 3213, 3616) clustered with ductal-origin samples (Figure 3A). These findings provide insights into the dual ontogeny of Progenitor:KP models.

**Figure 3.**
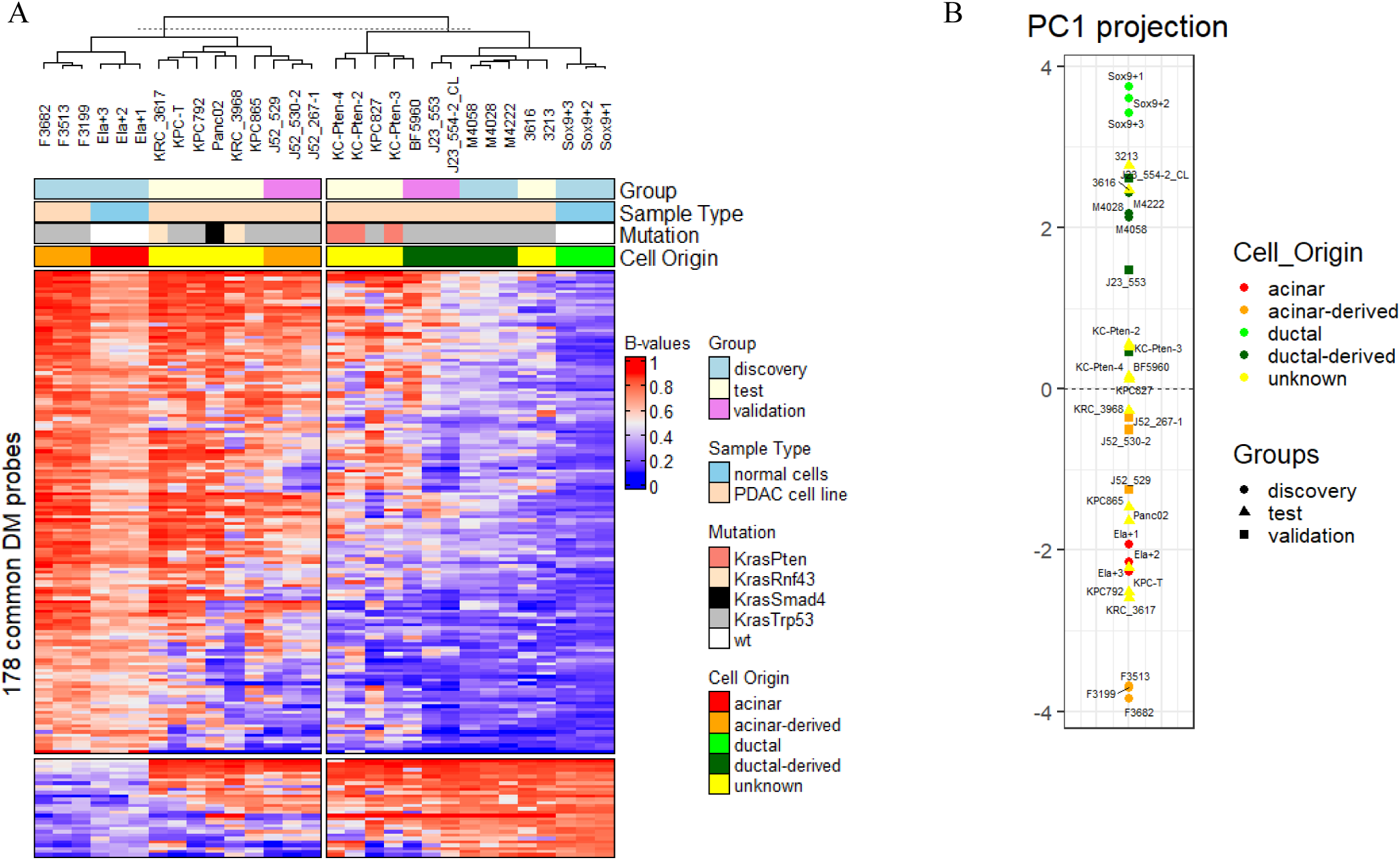
Progenitor:KP cell lines cluster based on their origin using the signature. (A) Hierarchical clustering of the discovery, validation and test samples using the 178 common probes. (B) Projection of the validation and test cell lines onto the PC1 derived from the discovery samples using the cell-of-origin signature.

We also explored the relationship between mutations found in intraductal papillary mucinous neoplasms (IPMN) and the cell of origin of PDAC. We profiled the DNA methylation of cell lines derived from mouse models developing IPMN, namely Ptf1a^Cre^; LSL- Kras^G12D/+^; LSL- Rnf43^fl/fl^ (Progenitor:KR, n = 2) [22] and Pdx1Cre; LSL- Kras^G12D/+^; LSL- Pten^fl/fl^ (Progenitor:KPten, n = 3) [23] mice. Using the cell-of-origin signature, progenitor:KR cell lines (KRC_3617, KRC_3968) clustered with the acinar-origin samples, while Progenitor:KPten cell lines (KC-Pten-2, KC-Pten-3, KC-Pten-4) clustered with the ductal- origin samples (Figure 3A). This suggests that the driver mutations not only influence the neoplastic fate (IPMN vs PanIN) but also the cellular origin of the tumor (acinar vs ductal).

Additionally, we analyzed the Panc02 cell line, which was derived by exposing the mice to a carcinogen, i.e. under non-genetically engineered conditions that more closely mimicthe sporadic PDAC formation in humans [24]. Despite the unknown cellular origin of the tumor arising after exposure to mutagenic agent, Panc02 cells clustered with the acinar-origin samples (Figure 3A).

To reinforce these results, the unknown cell-origin samples were projected onto the PCA space derived from the discovery dataset using the signature probes (Supplementary Figure 4A). The predicted cellular origins (Figure 3B, Supplementary Figure 4E) were consistent with those identified through hierarchical clustering (Figure 3A).

Overall, these findings underscore the effectiveness of the cell-of-origin DNA methylation signature in classifying pancreatic cancer cell lines, even when the cellular origin is unknown.

### Application of Logistic Regression Model for Cell-of-Origin Classification

To assess the predictive power of our cell-of-origin DNA methylation signature, we applied a logistic regression model to predict the cell of origin for both known and unknown PDAC samples. The model was trained on the discovery set, which included normal acinar (n = 3) and ductal cells (n = 3), as well as acinar-derived (n = 3) and ductal-derived PDAC cell lines (n = 3). The 178 common differentially methylated probes identified in our prior analysis served as features for this logistic regression model.

We first evaluated the model on the validation cell lines (n = 6), achieving an accuracy of 1.0 in predicting their cell of origin (Figure 4A). Classification was based on the probabilities generated by the model: a probability above 0.5 suggested a ductal origin, while a probability below 0.5 indicated an acinar origin. To address potential low-confidence predictions, we set a confidence threshold based on the lowest probability observed among the validation PDAC cell lines (0.436 for sample J52_267-1) (Supplementary Table S3). This threshold defined our confidence margins, such that samples with probabilities falling between 0.436 and 0.564 (1 - 0.4643) were considered low-confidence. These samples were not assigned a definitive origin and were instead considered as unclassified.

**Figure 4.**
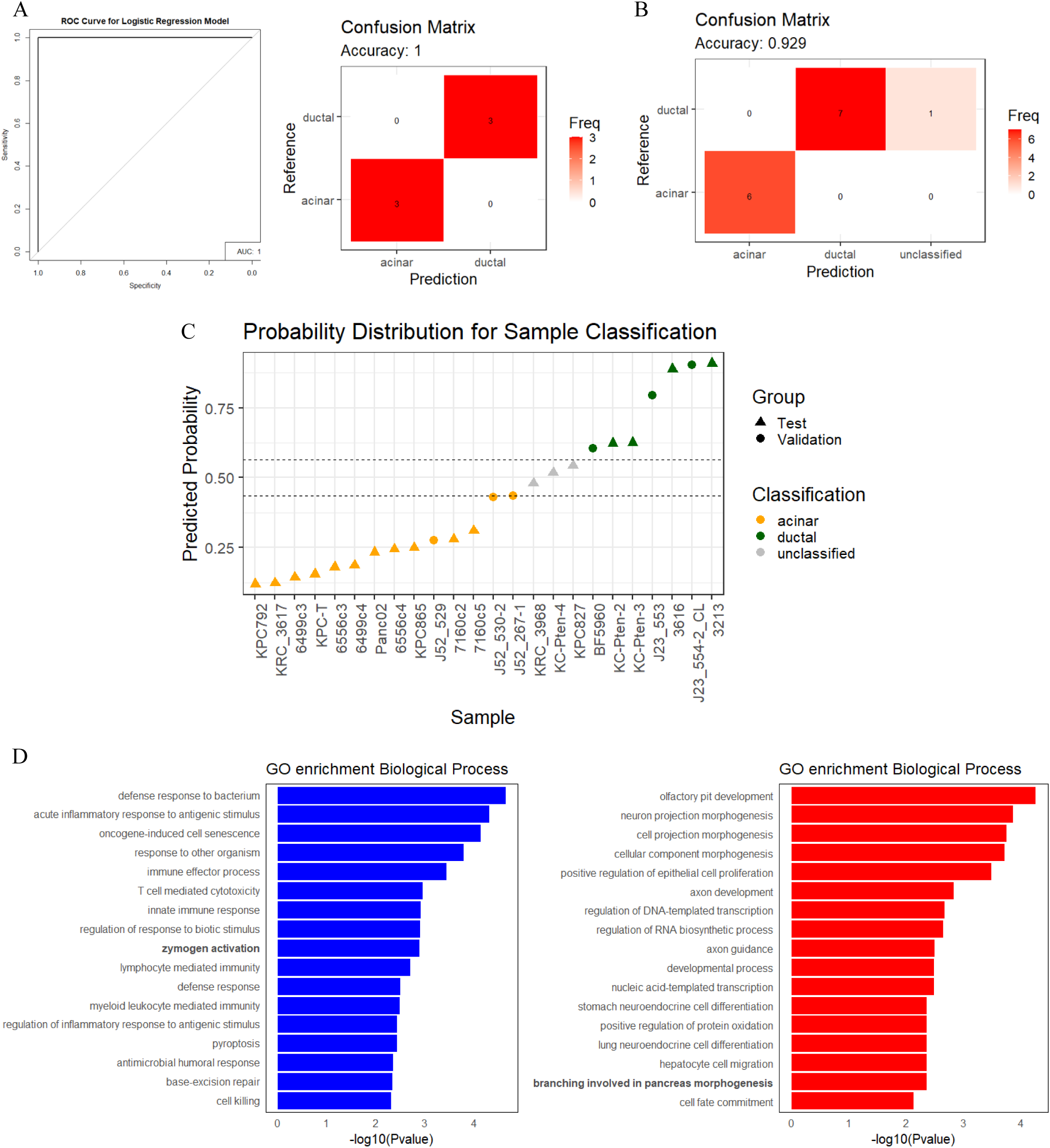
Predicting PDAC cell of origin, linked to distinct biological processes associated with acinar- and ductal-derived PDACs. (A) Receiver Operating Characteristic (ROC) curve and confusion matrix reporting the high accuracy (accuracy: 1) of the logistic regression model in predicting the cell of origin of the validation cell lines. AUC = area under the curve. (B) Confusion matrix reporting the high accuracy (accuracy: 0.929. Thirteen PDAC samples out of 14 correctly assigned) of the logistic regression model in predicting the cell of origin of the validation samples. (C) Distribution plot of predicted probabilities extracted from the logistic regression model. Samples with probability lower than the threshold are classified as acinar (orange). Samples with probability higher than the threshold are classified as ductal (green). Samples with probabilities falling within the threshold margins (dashed lines) are not classified and are shown in grey. The threshold is set as the lowest probability among the validation PDAC cell lines (0.436 for J52_267-1), and the threshold margins are defined as this threshold and (1-threshold). (D) Gene Ontology (GO) enrichment analysis of the 100 top- ranked differentially methylated promoters in acinar-derived (n = 11) vs ductal-derived (n = 10) PDAC cell lines, as classified in Table 1. Blue bars represent biological process (BP) ontology terms enriched in hypomethylated acinar-derived promoters and red bars represent BP ontology terms enriched in hypomethylated ductal-derived promoters. The analysis was performed using the performGoEnrichment.diffMeth() function of the RnBeads package in R software with default settings. Significant enrichment was considered with p-value < 0.01.

**Table 1.** Table providing an overview of the cell lines used in the study, including their classification based on their cell of origin as predicted by hierarchical clustering or logistic regression model. Cell lines in bold show assignment that differs between the two approaches.

| Cell line | Cell Origin | Group | Hierarchical clustering<br>Supplementary Fig.5 | Logistic regression |  |
| --- | --- | --- | --- | --- | --- |
| F3199 | acinar-derived | discovery | acinar | acinar | Acinar1:KP |
| F3513 | acinar-derived | discovery | acinar | acinar | Acinar1:KP |
| F3682 | acinar-derived | discovery | acinar | acinar | Acinar1:KP |
| M4028 | ductal-derived | discovery | ductal | ductal | Ductal1:KP |
| M4058 | ductal-derived | discovery | ductal | ductal | Ductal1:KP |
| M4222 | ductal-derived | discovery | ductal | ductal | Ductal1:KP |
| J52_529 | acinar-derived | validation | acinar | acinar | Acinar2:KP |
| J52_530-2 | acinar-derived | validation | acinar | acinar | Acinar2:KP |
| J52_267-1 | acinar-derived | validation | acinar | acinar | Acinar2:KP |
| J23_553 | ductal-derived | validation | ductal | ductal | Ductal2:KP |
| J23_554-2_CL | ductal-derived | validation | ductal | ductal | Ductal2:KP |
| BF5960 | ductal-derived | validation | ductal | ductal | Ductal2:KP |
| KPC792 | unknown | test | acinar | acinar | Progenitor:KP |
| **KPC827** | unknown | test | ductal | unclassified | Progenitor:KP |
| KPC865 | unknown | test | acinar | acinar | Progenitor:KP |
| 3213 | unknown | test | ductal | ductal | Progenitor:KP |
| 3616 | unknown | test | ductal | ductal | Progenitor:KP |
| 6499c3 | unknown | test | acinar | acinar | Progenitor:KP |
| 6499c4 | unknown | test | acinar | acinar | Progenitor:KP |
| 6556c3 | unknown | test | acinar | acinar | Progenitor:KP |
| 6556c4 | unknown | test | acinar | acinar | Progenitor:KP |
| 7160c2 | unknown | test | acinar | acinar | Progenitor:KP |
| 7160c5 | unknown | test | acinar | acinar | Progenitor:KP |
| KPC-T | unknown | test | acinar | acinar | Progenitor:KP |
| KC-Pten-2 | unknown | test | ductal | ductal | Progenitor:KPTen |
| KC-Pten-3 | unknown | test | ductal | ductal | Progenitor:KPten |
| **KC-Pten-4** | unknown | test | ductal | unclassified | Progenitor:KPten |
| KRC_3617 | unknown | test | acinar | acinar | Progenitor:KR |
| **KRC_3968** | unknown | test | acinar | unclassified | Progenitor:KR |
| Panc02 | unknown | test | acinar | acinar | Panc02 |

Next, we included primary tumors (n = 5) and orthotopic allografts (n = 3) of the validation group. Using the trained model with the cell-of-origin DNA methylation signature, we accurately classified these tumors, achieving an overall accuracy of 0.929 (Figure 4B). This high accuracy underscores the robustness and distinctiveness of the DNA methylation patterns captured by our signature, supporting their utility as reliable biomarkers for cell-of-origin classification.

We also applied the classifier to the test set (n = 12). The model successfully predicted the cell of origin for 9 out of 12 progenitor-derived cell lines (Figure 4C). These predictions were consistent with the hierarchical clustering results (Table 1). High-confidence predictions were made for five Progenitor:KP cell lines (KPC792, probability = 0.119; KRC_3617, probability = 0.124; KPC-T, probability = 0.155; 3213, probability = 0.908; 3616, probability = 0.889). However, three cell lines (KRC_3968, KC-Pten-4 and KPC827) were not classified due to low-confidence predictions (Figure 4C, Table 1).

Finally, we sought to determine whether tumor cells derived from the same mouse but exhibiting different characteristics correspond to different cell origins. To investigate this, we analyzed pairs of cancer cell clones generated from three Progenitor:KP mice (n = 6) [25]. When grafted into immunocompetent mice, the clones of each pair produced tumors with either high or low T cell infiltration, reflecting distinct epigenetic and transcriptomic profiles (Table 2) [25]. Applying the classifier to these cell lines revealed that both clones in each pair were consistently assigned the same origin (acinar origin), regardless of their diverse immune properties (Figure 4C, Table 1). The predictions matched the hierarchical clustering results based on the cell-of-origin signature, showing that each pair clustered together and confirming their close proximity (Supplementary Figure 5A).

**Table 2.** Table providing information about the T-cell infiltration status of pancreatic cancer cell clones derived from Progenitor:KP mice [25].

| **Cell line** | **T-cell infiltration status** | **Pair** |
| --- | --- | --- |
| 6499c3 | Low | 1 |
| 6499c4 | High | 1 |
| 6556c3 | Low | 2 |
| 6556c4 | High | 2 |
| 7160c2 | High | 3 |
| 7160c5 | Intermediate | 3 |

In conclusion, the logistic regression model based on our DNA methylation signature provides a powerful tool for classifying pancreatic tumor samples by their cell of origin. This approach not only confirms the validity of our signature but also offers a practical method for predicting cell-of-origin in PDAC models with unknown cellular origins.

### Distinct Characteristics of Acinar-Derived and Ductal-Derived PDAC

Based on the logistic regression classification, we performed differential methylation analysis comparing the 11 acinar-origin to the 11 ductal-origin cell lines. To understand the biological significance of the DNA methylation profiles specific to acinar-derived and ductal-derived cancer cells, we focused on the differentially methylated promoters, as methylated promoter regions are usually associated with transcriptional repression [26].

We first identified the top-ranked hypomethylated promoters in acinar-derived (n = 45) and ductal-derived (n = 76) PDAC cell lines and then performed Gene Ontology (GO) enrichment analysis using the performGoEnrichment.diffMeth() function of the RnBeads package in R software with default settings. The top ontology terms enriched in hypomethylated promoters in acinar-derived PDAC cell lines included biological processes linked to immune and inflammatory response (Figure 4D). Interestingly, the promoters of *Prss3b* (aka *2210010C04Rik*) (mean difference = -0.192, false discovery rate (FDR)-adjusted combined p-value = 0.061), *Klk1* (mean difference = -0.305, comb.p.adj.fdr = 0.032) and *Nlrp6* (mean difference = -0.288, comb.p.adj.fdr = 0.004) genes, which are enriched for the term zymogen activation, were hypomethylated in acinar-derived cells (Supplementary Figure 6A, Supplementary Table S4).

On the other hand, hypomethylated promoters in ductal-derived cells displayed enrichment for GO terms related to transcriptional regulation, morphogenesis, cell fate and differentiation (Figure 4D). Notably, the branching involved in pancreas morphogenesis process, which shape the ductal network, was enriched in ductal-derived cells (Figure 4D, Supplementary Table S4). These findings demonstrate that the molecular profile of pancreatic tumors is dependent on the cell of origin and indicate that some cell-specific biological functions are epigenetically imprinted.

Additionally, the promoter of *Khdrbs3* (mean difference = 0.406, comb.p.adj.fdr = 0.009), which is linked with multi-drug resistance [27–29] was significantly hypomethylated in ductal-derived PDAC cells (Supplementary Figure 6B). Given this, we aimed to determine the sensitivity of acinar-derived and ductal-derived PDAC cell lines to common chemotherapeutic agents. We assessed cell survival after 72-hour treatment with gemcitabine (GEM) and 5-fluorouracil (5-FU), using proliferation assays. Acinar1:KP (n = 3) and Ductal1:KP (n = 3) cancer cells responded differentially to the tested drugs, with ductal-derived cells being significantly more resilient (GEM tolerance, p-value = 0.022; 5-FU tolerance, p-value = 0.0002, ANCOVA) (Figure 5A). The colony-forming ability (Figure 5B- C) and migratory capacity (Figure 5D-E) of acinar-derived cells after drug treatment were highly impacted compared to ductal-derived cells, confirming the results of the drug sensitivity assay. Overall, these results suggest that the cell-of-origin correlates with distinct drug responses, reinforcing the potential of being used for therapeutic stratification in pancreatic cancer.

**Figure 5.**
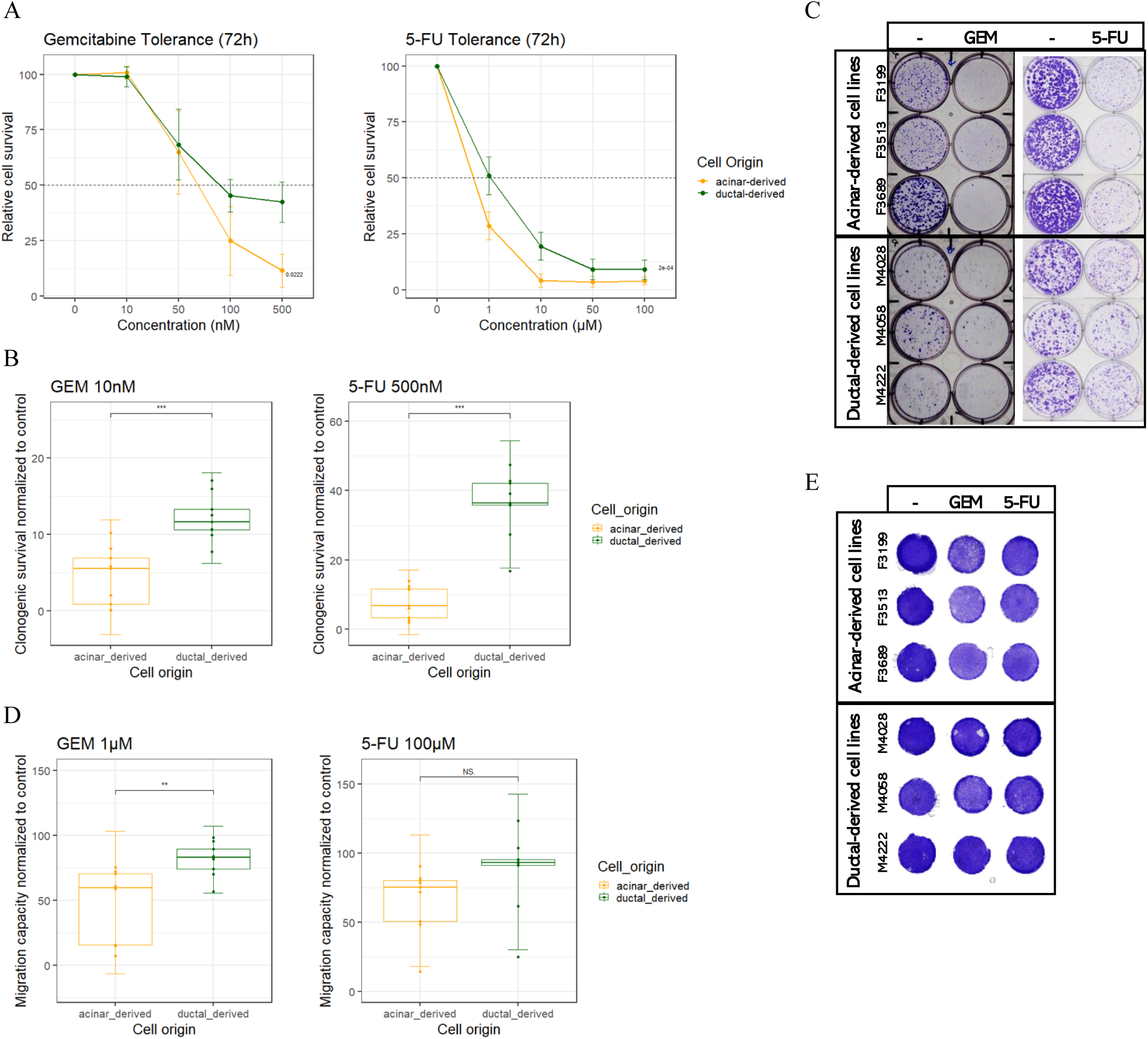
Acinar-derived PDAC cell lines respond better to chemotherapy agents. (A) GEM and 5-FU tolerance of acinar-derived (orange, n=3) and ductal-derived (green, n=3) cell lines was assessed using a proliferation assay after 72 hours of treatment with varying concentrations of the drugs. The horizontal dashed line indicates a relative cell survival of 50%. The experiment was performed in triplicate. Data are expressed as mean ± standard deviation, and statistical significance was determined using Analysis of Covariance (ANCOVA). (B) The colony- forming ability of acinar-derived (orange, n=3) and ductal-derived (green, n=3) cell lines was assessed after 7 days of treatment with 10 nM GEM and 500 nM 5-FU. The experiment was performed in triplicate. Data are expressed as mean ± standard deviation, and statistical significance was determined using Student’s t-test. (C) Representative images of the clonogenic assay performed on acinar-derived (F3199, F3513, F3682) and ductal-derived (M4028, M4058, M4222) PDAC cell lines. (D) The migratory capacity of acinar-derived (orange, n=3) and ductal-derived (green, n=3) cell lines was assessed using Transwell inserts (8 µm pore size) after 24 hours of treatment with 1µM GEM or 100µM 5-FU. The experiment was performed in triplicate. Data are expressed as mean ± standard deviation, and statistical significance was determined using Student’s t-test. (E) Representative images of the membranes of the Transwell inserts, showing non-migratory cells from the acinar-derived (F3199, F3513, F3682) and ductal-derived (M4028, M4058, M4222) PDAC cell lines.

Finally, the promoter of *Krt20* (mean difference = -0.282, comb.p.adj.fdr = 0.007), a marker associated with the classical PDAC subtype [5] and known to be overexpressed in acinar-derived PDACs [9], was significantly hypomethylated in acinar-derived PDAC cells (Supplementary Figure 6C). Interestingly, KRT20 was expressed in 5/35 PDAC [30], contrary to other Keratins (KRT7, KRT18, KRT19) which were expressed in 100% of PDAC. This suggests an association between cell of origin and PDAC heterogeneity.

## Discussion

Numerous studies using mouse models have established a strong association between PDAC’s ontogeny and its heterogeneity [7–10,13]. Although the concept of determining the cell of origin as a decisive factor for subclassifying tumors is gaining traction, there is no applicable method to trace back the lineage of pancreatic tumors. Here, we address this need by developing a DNA methylation-based classification system that leverages the well-known retention of cell-specific epigenetic imprints in tumors [17,18,31,32]. Using comprehensive DNA methylation profiling of normal pancreatic acinar and ductal cells, as well as their corresponding PDAC cell lines, we identified persistent DNA methylation patterns linked to the cellular origin of PDAC. These patterns formed the basis of a robust cell-of-origin DNA methylation signature, which reliably distinguished acinar-derived from ductal-derived PDACs across diverse experimental settings.

Our findings revealed distinct methylation profiles in normal acinar and ductal cells, with significant differences observed at thousands of CpG sites. These results are consistent with prior studies of human pancreatic cells [12,14]. Interestingly, we observed that PDAC cell lines derived from acinar (Acinar1:KP) and ductal (Ductal1:KP) origins exhibited partial loss of their original methylation patterns. This partial erasure of cell identity is consistent with established concepts of epigenetic reprogramming during carcinogenesis [12,15,16,33] and may also reflect the effects of cell culture [14]. Despite these changes, lineage-specific methylation markers were retained, forming a distinct signature that reliably traced the cellular origin of tumors. This finding reinforces earlier observations that tumors from different lineages maintain distinct epigenetic markers [17,18,31,32].

To evaluate the sensitivity of our cell-of-origin DNA methylation signature, we analyzed independent acinar-derived (Acinar2:KP) and ductal-derived (Ductal2:KP) PDAC cell lines. Unexpectedly, global methylation profiling showed that Acinar2:KP samples clustered closer to ductal-derived PDAC cells than to Acinar1:KP. This discrepancy may reflect differences in tumor generation protocols and culturing conditions. For example, Acinar1:KP tumors were induced by cerulein treatment, which is known to cause sustained transcriptional and epigenetic reprogramming in acinar cells [34]. Also, Acinar2:KP samples were derived from mice with haploinsufficiency for *Ptf1a*, as the construct used to express CreER was inserted into the *Ptf1a* gene. Ptf1a is a transcription factor essential to maintain the acinar cell fate. Consequently, a decreased expression contributes to acinar-ductal metaplasia [35] and affects how PDAC is initiated, compared to the mice used to generate the Acinar1:KP samples [36]. Nevertheless, when assessed with our methylation signature, Acinar2:KP cells clustered correctly with acinar-derived samples, highlighting the signature’s ability to accurately capture cell-of-origin characteristics.

Validation using orthotopic allografts and primary tumors further demonstrated the robustness and reproducibility of the cell-of-origin signature. The signature consistently distinguished acinar- and ductal-origin tumors even in the presence of heterogeneous stroma [37,38], adding a new layer of reliability. These results align with earlier studies demonstrating the stability of lineage-specific methylation among various cancer models [17,33].

The progenitor-derived PDAC models have been in the scientific spotlight for decades, as it is considered the most relevant preclinical model for studying PDAC, due to the significant heterogeneity of its tumors, similar to what is observed in humans. Numerous studies have attempted to explain this heterogeneity, which is characterized by the simultaneous presence of several cancer cell states which evolve dynamically during tumor progression [20,39]. In these models, oncogenic mutations are introduced at the embryonic stage, leaving the cellular origin of the resulting tumors unclear. To address this, we analyzed progenitor-derived tumors and provided new insights into their ontogeny. Using our cell-of-origin DNA methylation signature, we classified Progenitor:KP (aka KPC) cell lines [20–22,25], which express mutated *Kras* and *Trp53*. Hierarchical clustering revealed that the majority of the analyzed cells were of acinar-origin, while the remainder were of ductal-origin. These results suggest that KPC tumors can originate from two different cell types, and that the primary factor driving tumor heterogeneity in this model is the cell of origin, although cellular plasticity may further contribute to this diversity. This raises important questions about previous studies using KPC mice. If tumors in this model have different cellular origins, it could impact the interpretation of drug treatment outcomes. Translating this concept to humans suggests that human PDAC may also originate from different cell types, with potential clinical implications. For instance, drug resistance could result from the presence of cancer cells with different origins, each responding differently to the same treatment. Developing a strategy to identify the cell of origin in human PDAC could not only deepen our understanding of tumor development but also help design more personalized and effective therapies.

We further explored the relationship between PDAC pathophysiology and cellular origin by profiling cell lines derived from GEMMs with mutations associated with IPMN formation. IPMNs, precancerous cystic lesions, were initially thought to arise from ductal cells [40–43]. However, acinar cells harboring mutations in *Gnas*, a key IPMN-associated oncogene in humans [44], can also give rise to these lesions [45]. Our analysis showed that Progenitor:KR cell lines with *Rnf43* loss [22], a gene frequently mutated in human IPMNs [46], clustered with acinar-origin samples, whereas Progenitor:KPten cell lines [23], with *Pten* ablation, clustered with ductal-origin samples. These findings align with prior observations that only ductal-specific *Pten* loss leads to IPMN-mediated PDAC formation [42], but also confirm that IPMN can derive from acinar cells. Overall, our results from progenitor-derived PDAC GEMMs emphasize the interplay between driver mutations and cell type in determining neoplastic outcomes.

The 178 CpG probes constituting our cell-of-origin signature were resilient enough to serve as reliable biomarkers for lineage classification. A logistic regression classifier trained on this signature achieved high accuracy in distinguishing acinar- and ductal-origin PDACs, including primary tumors with complex microenvironments. Furthermore, the classifier successfully predicted the cell of origin for previously unclassified PDAC samples from progenitor-derived models, with predictions aligning closely with hierarchical clustering results. This robust performance provides a promising foundation for potential application in clinical diagnostics. We tried to apply our signature in human samples, but only around tweenty probes were syntenic between the two species, which was not a sufficient number to carry out a correct analysis. Nevertheless, we believe that a similar approach could lead to the identification of a human signature.

The biological relevance of cellular origin was further demonstrated by differences in methylation patterns linked to tumor behavior. Acinar-derived PDAC cells exhibited hypomethylation of immune- and inflammatory-response-associated promoters, consistent with their immunogenic profile [13]. On the other hand, ductal-derived PDAC cells displayed hypomethylation of genes associated with migration and proliferation, consistent with their more aggressive phenotype [9,10,12]. Additionally, the hypomethylation of *Khdrbs3*, linked to multidrug resistance [27–29], provides an epigenetic explanation for the chemoresistance observed in ductal-derived PDAC cells, a finding that may inform treatment strategies based on tumor origin.

In conclusion, our findings underscore the significance of DNA methylation as both a marker of cellular origin and a modulator of tumor behavior in PDAC. Future studies should validate this approach in human PDAC samples and evaluate its clinical potential for patient stratification and therapeutic targeting. The preservation of cell-specific methylation signatures offers novel insights into PDAC biology and holds promise for advancing diagnostics and treatment for this lethal cancer.

## Acknowledgements

The authors thank the members of the LPAD laboratory, Frédéric Lemaigre for discussions, Bruno Maricq for help in mouse care, Marine Fellmann for technical help, and Sebastian Kobold, Jennifer Morton, Anirban Maitra for cell lines.

## Ethics Statement

This study does not involve human participants. All experimental procedures performed on mice were approved by the animal welfare committee of the University of Louvain Medical School (reference 2021/UCL/MD/054).

---

**Supplementary Figure 1.**
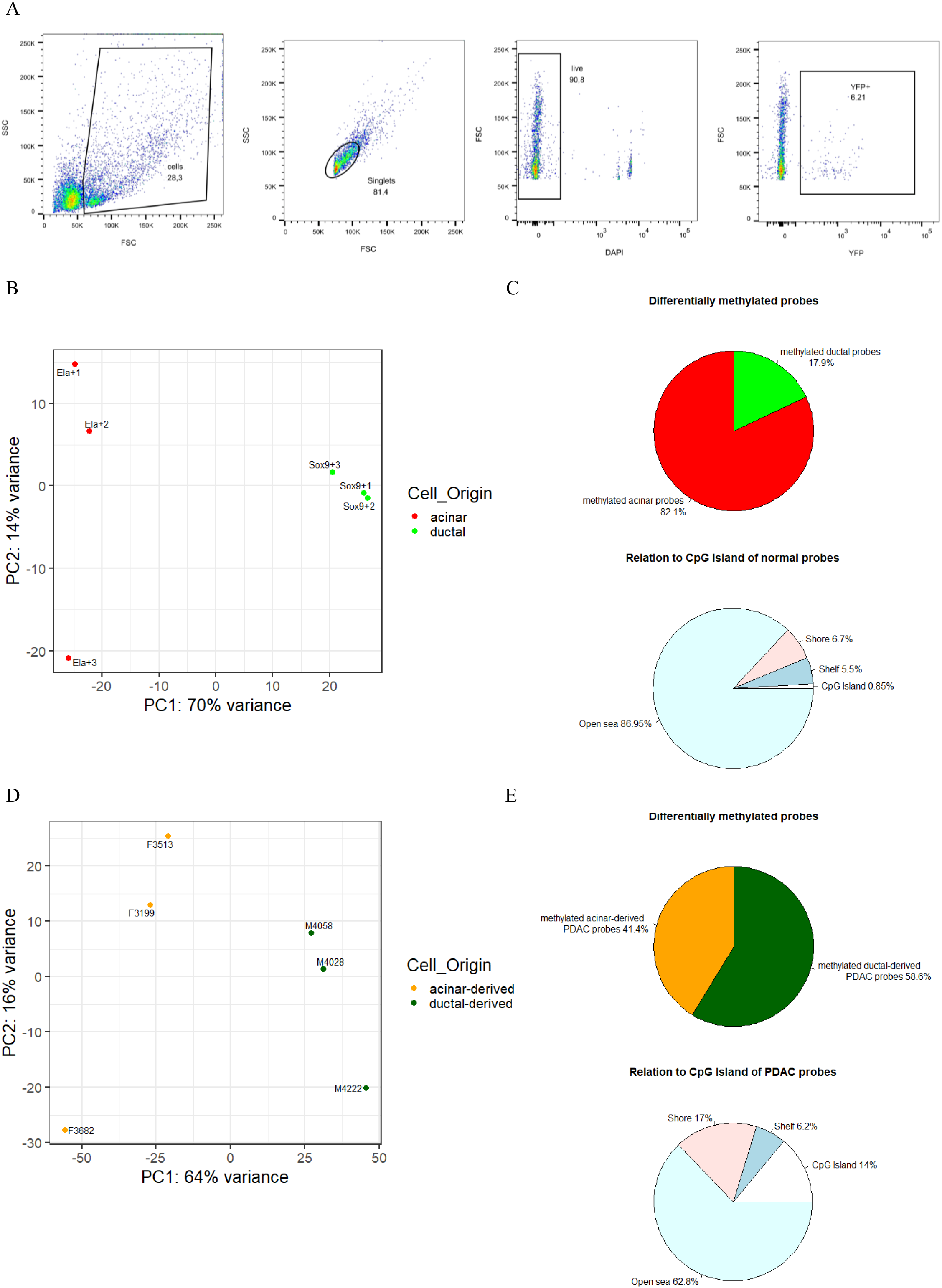
(A) Representative flow cytometry plots illustrating the FACS sorting strategy to isolate acinar cells (YFP+. The SSC channel sorts cells based on their granularity (i.e intracellular complexity). The numbers in the graphs indicates the percentage of gated cells. (B) PCA of the methylation levels from normal FACS-purified acinar and ductal cells. Percentages indicate the proportion of variance explained by each component. Acinar (Acinar:YPF): n = 3; Ductal (Ductal:EGFP): n = 3. (C) Distribution of the 2,346 normal DMPs with reference to cell origin (top) and genomic features (bottom). CpG island are DNA regions frequently found in gene promoters and rich in CpG sites, compared to the rest of the genome. Based on their proximity to CpG islands, other DNA regions are categorized as shores, shelves, and open sea, with shores being the closest, and the open sea the most distant from CpG islands. (D) PCA of the methylation levels from the acinar-derived and ductal-derived PDAC cell lines. Percentages indicate the proportion of variance explained by each component. Acinar-derived (Acinar1:KP): n = 3; Ductal-derived (Ductal1:KP): n = 3. (E) Distribution of the 6,135 PDAC DMPs with reference to cell origin (top) and genomic features (bottom).

**Supplementary Figure 2.**
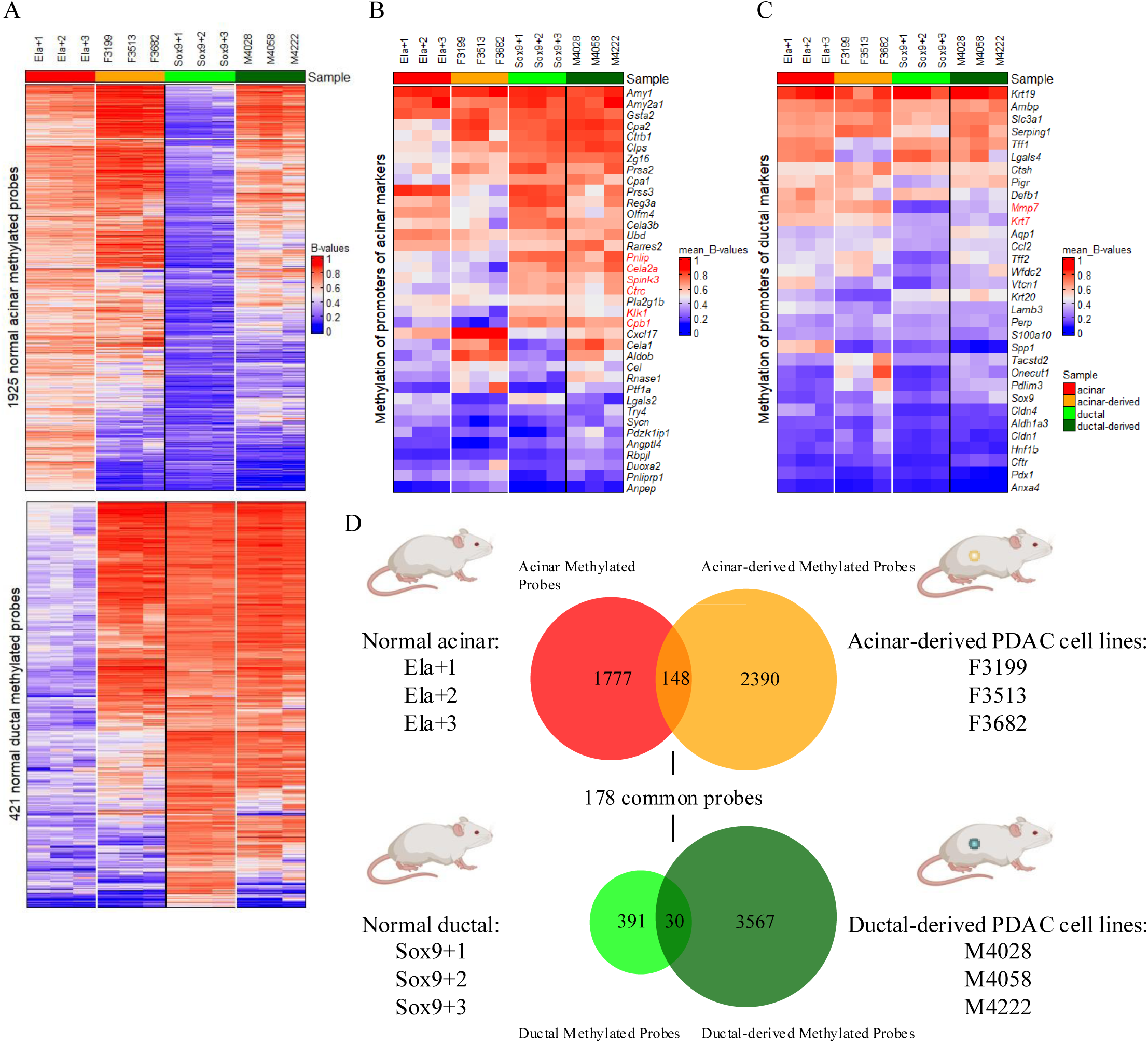
(A) DNA methylation status in acinar, ductal, acinar-derived, and ductal-derived samples of the most highly methylated acinar (top) and ductal (bottom) probes. ComplexHeatmap was used to generate the plots, and image rasterization (raster_quality = 10) was used when features exceeded 2,000. (B) Mean methylation levels of probes located in the promoters of acinar markers, in acinar and ductal cells, and acinar-derived, and ductal-derived cell lines. Acinar-specific genes that retained their promoter methylation profiles in acinar cells and acinar-derived cell lines are highlighted in red. (C) Mean methylation levels of probes located in the promoters of ductal markers, in acinar and ductal cells, and acinar-derived, and ductal-derived cell lines. Ductal-specific genes that retained their promoter methylation profiles in acinar cells and acinar-derived cell lines are highlighted in red. (D) Scheme illustrating the composition of the discovery group and the strategy followed to identify the 178 common probes used as the cell-of-origin DNA methylation signature in the study.

**Supplementary Figure 3.**
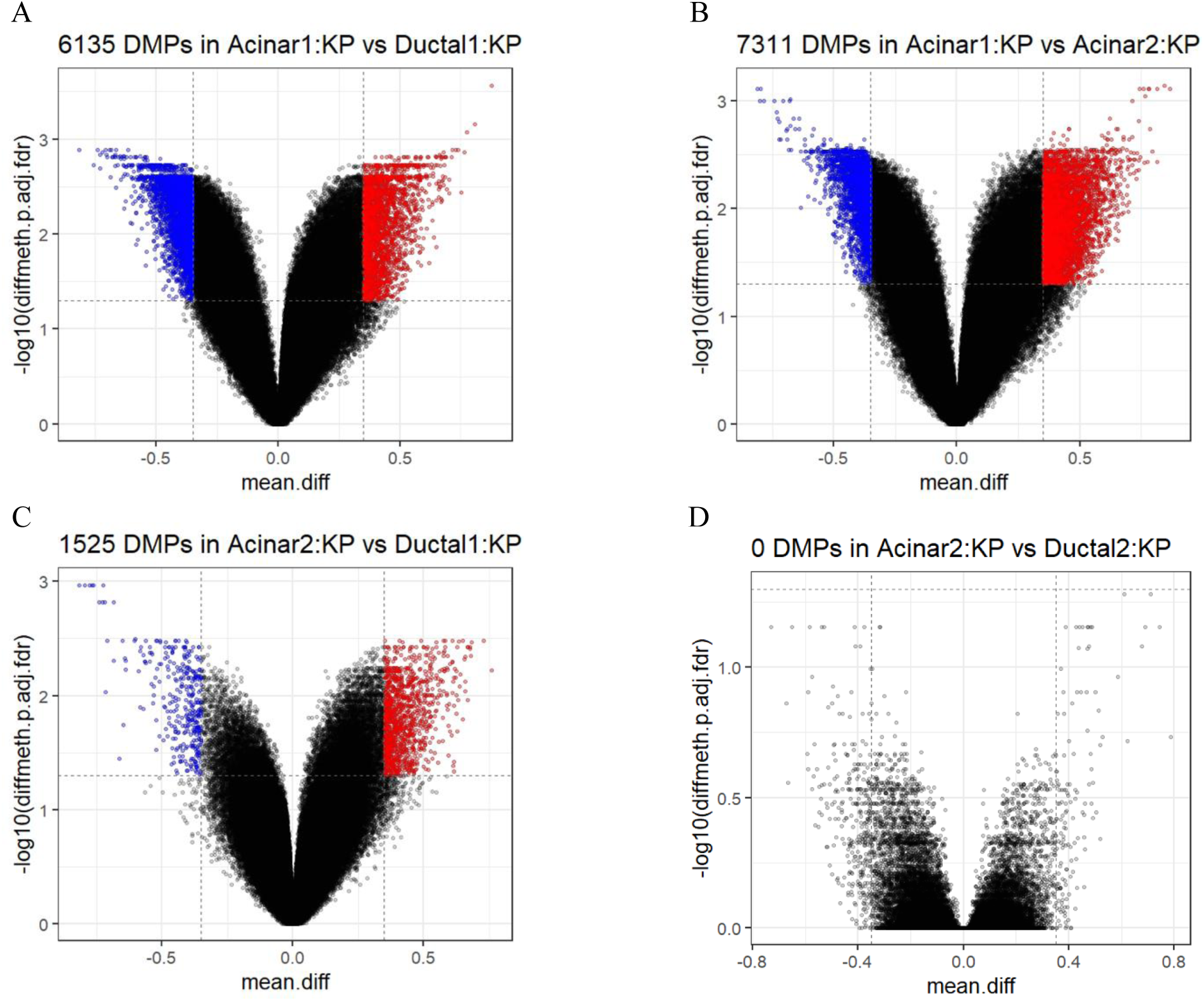
Volcano plot of top differentially methylated probes in Acinar1:KP vs Ductal1:KP (A), Acinar1:KP vs Acinar2:KP (B), Acinar2:KP vs Ductal1:KP (C) and Acinar2:KP vs Ductal2:KP cell lines (D). Vertical dashed lines indicate an absolute mean difference of 0.35. Horizontal dashed line corresponds to an FDR adjusted p-value equal to 0.05. Red dots represent hypermethylated probes and blue dots represent hypomethylated probes.

**Supplementary Figure 4.**
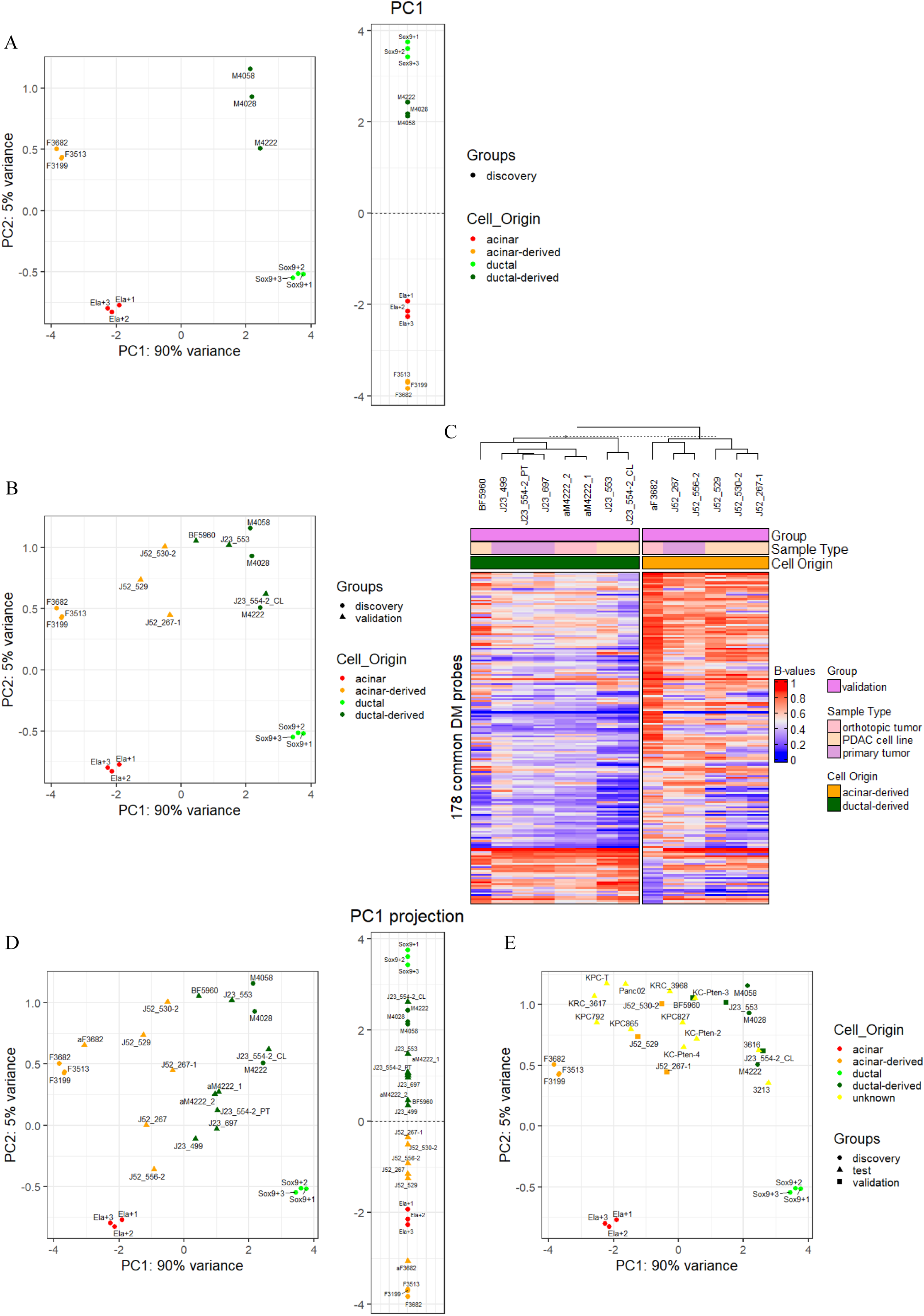
(A) PCA (left) of the discovery samples and their distribution according to PC1 (right) using the cell-of-origin signature. (B) Projection of validation cell lines onto the PCA space derived from the discovery samples using the cell-of-origin signature. (C) Hierarchical clustering of the validation samples using the cell-of-origin signature. (D) Projection of validation samples onto the PCA space (left) and PC1 (right) derived from the discovery samples using the cell-of-origin signature. (E) Projection of validation and test samples onto the PCA space derived from the discovery samples using the cell-of-origin signature.

**Supplementary Figure 5.**
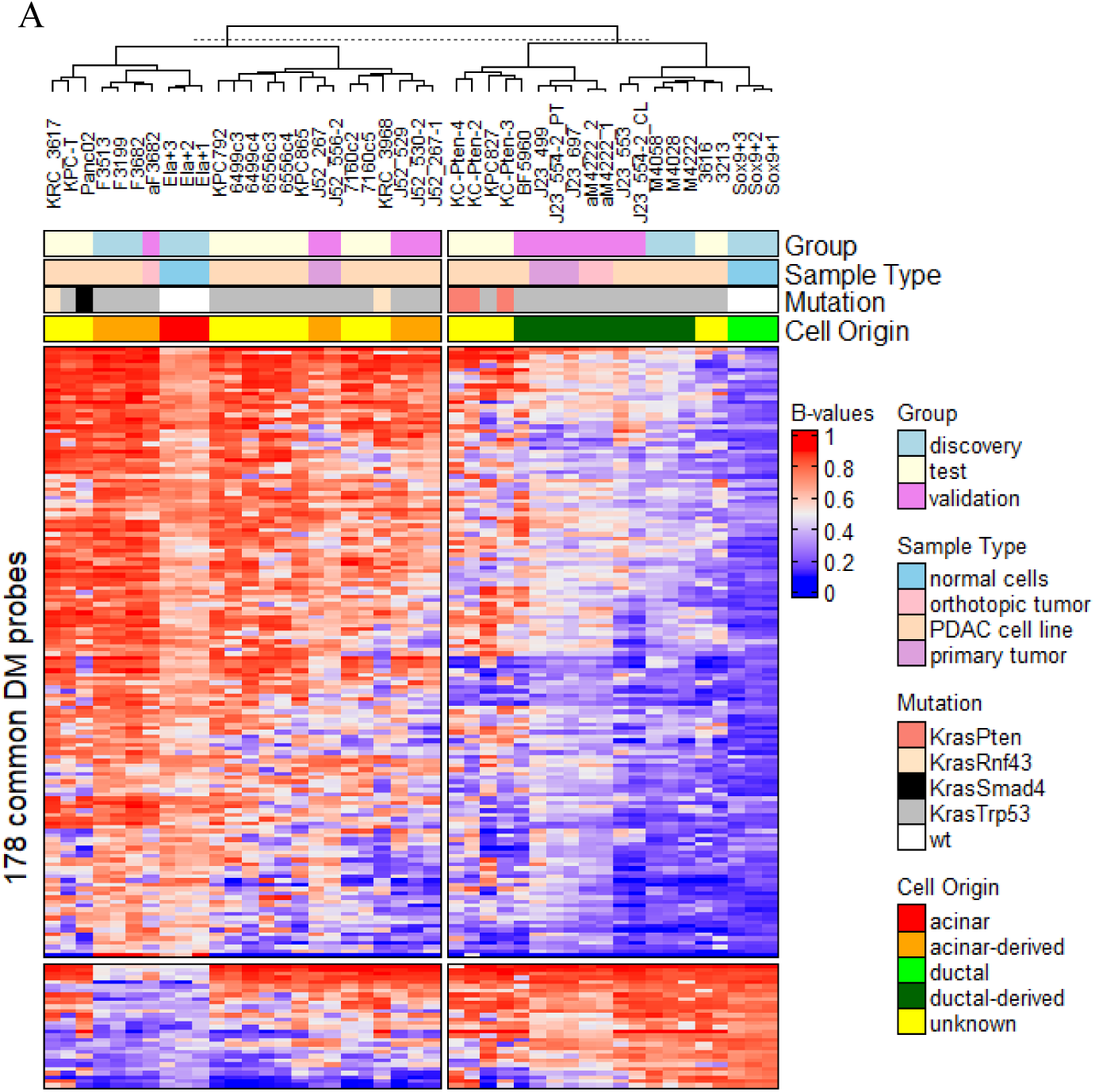
(A) Hierarchical clustering of the 44 samples used on this study using the cell-of-origin signature, including the 3 pairs of Progenitor:KP cell lines with different immune properties.

**Supplementary Figure 6.**
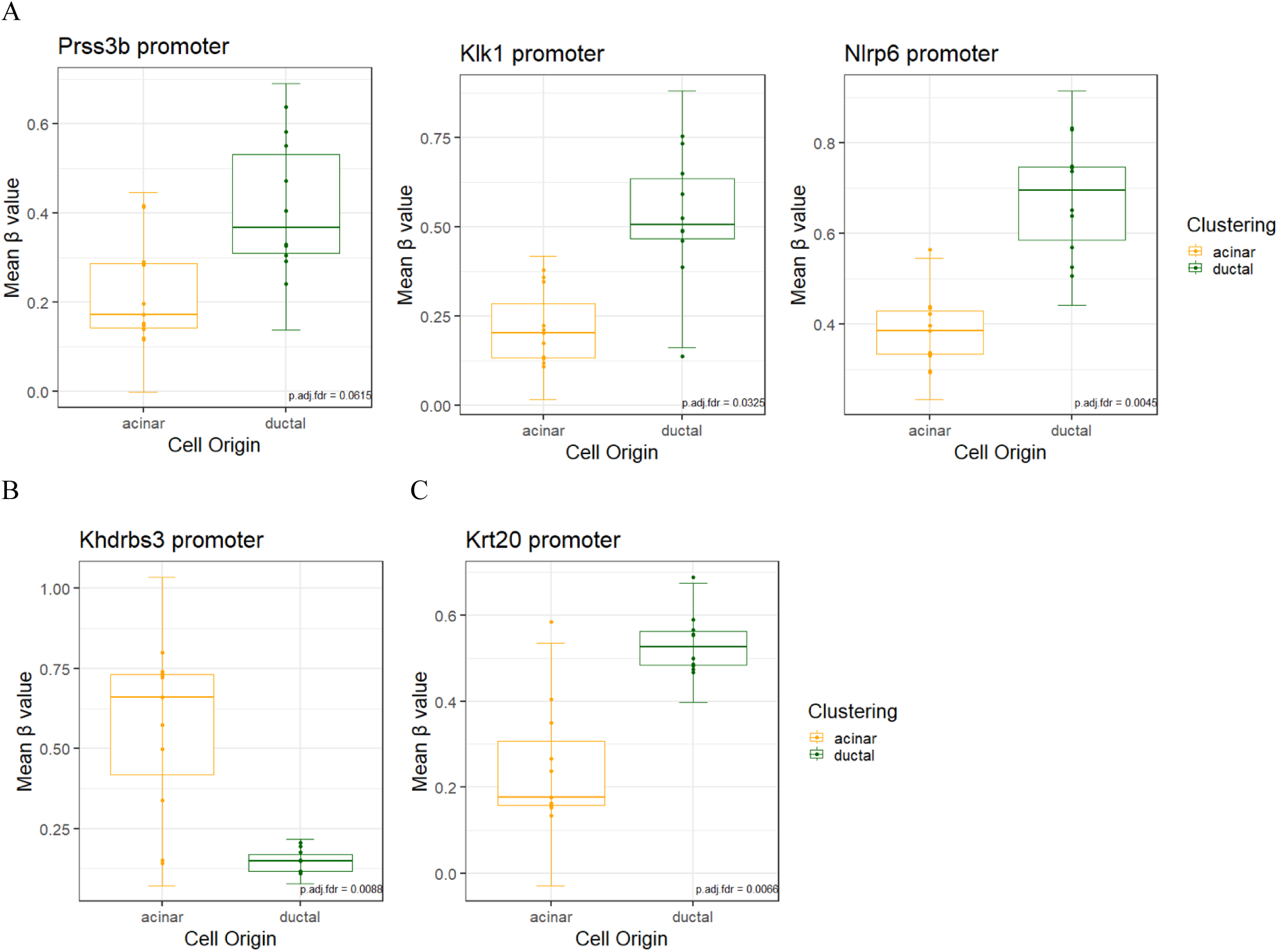
(A) Boxplots of the mean methylation of gene promoters enriched for GO:0031638 term (zymogen activation) in acinar-derived (orange, n=11) vs ductal-derived (green, n=10) PDAC cell lines as classified in Table 1. Boxplot of the mean methylation of *Khdrbs3* (B) and *Krt20* (C) gene promoter in acinar-derived (orange, n=11) and ductal-derived (green, n=10) PDAC cell lines as classified in Table 1. (A-C) Lines represent the mean ± standard deviation. The false discovery rate (FDR)-adjusted combined p-value, using a generalization of Fisher’s method, is plotted in the lower right corner.

